# Entropy regularised reinforcement learning reconciles aversive and action prediction errors in the tail of the striatum

**DOI:** 10.64898/2026.07.09.737461

**Authors:** Pranav Mahajan, Ben Seymour

## Abstract

Dopamine activity in the tail of the striatum (TS) presents a novel challenge for reinforcement-learning theories of dopamine. Some studies suggest that TS-projecting dopamine signals encode aversive or threat prediction errors, whereas others argue that they encode action prediction errors involved in soft-habit formation. Here, we show that these accounts need not be mutually exclusive. We instantiate an entropy-regularised reinforcement-learning model in which TS-projecting dopamine neurons update both aversive values and the default policy. In this model, threat belief gates aversive value initialisations, producing TS-like activity during retreat from potentially threatening novel objects, while default-policy learning generates action prediction error signals that decline as actions become habitual. Our results further suggest why both of these signals may need to coexist in the temporal difference errors in our model, and qualitatively reproduce key response patterns and simulations from studies previously used to support both views. Beyond this descriptive reconciliation, our model simulations also highlight the normative role of the tail of the striatum in cautious behaviours in the context of potential threats and stable learning in the face of outcome uncertainty.

## 1 Introduction

The proposal that phasic activity in midbrain dopamine (DA) neurons encodes temporal-difference reward prediction errors has provided a foundational computational account of dopamine function [1]. By leveraging the temporal difference (TD) learning algorithm [2], this account suggested that the brain could assign credit using temporally successive predictions of expected future reward. Numerous experiments have since supported this relationship in canonical reward-learning settings within the TD reinforcement learning (TDRL) framework [3, 4, 5, 6, 7].

A core feature of the Schultz et al. [1] model is the scalar, globally broadcast reward prediction error (RPE) signal in DA responses. This is not a superficial claim but has direct implications for decision-making. For instance, the TDRL framework allows agents to learn and compare the values (i.e., estimates of long-run potential outcomes) for different states and actions, enabling agents to take better actions. Early physiological reports also highlighted this homogeneity of responses of midbrain DA neurons on simple conditioning tasks, especially that most units respond to unexpected reward [8].

However, a body of recent work demonstrates remarkable heterogeneity in these midbrain dopamine responses, tuned to different rewards [9, 10, 11, 12, 13], punishments [14, 15, 16, 17, 18], and sensorimotor variables [19, 20, 21, 22, 23, 24, 25, 26]. These DA responses and agent behaviour are further significantly shaped by the environmental context and belief states [27, 28, 13]. While action selection ultimately requires an integration of multiple value signals, the notion of a monolithic RPE signal is increasingly at odds with observations of diverse dopamine responses, both between and within striatal subregions.

Some between-target heterogeneity, particularly responses specific to different reward or punishment types, can in principle be captured by extending the classical model [1] to multiple rewarding objectives where different reward-specific values can be recombined to drive adaptive behaviour [29]. However, this still leaves heterogeneity within DA targets [30], D1 and D2 pathways [31, 18], novelty-based bonuses [32, 17], and action-specific responses [26] unaccounted for.

Nowhere is this challenge more apparent than in the tail of the striatum (TS). Currently, there is considerable debate over whether TS-projecting dopamine encodes action prediction errors [26] or aversive/threat prediction errors [15, 16, 17]. On one hand, evidence suggests TS neurons encode threat predictions that are updated by TS-projecting threat prediction errors in the substantia nigra lateralis (SNl) [16]. TS dopamine responds not only to aversive outcomes (e.g., airpuffs) [15, 16], but also during retreat from potentially threatening novel objects, and shapes avoidance behaviours in animals [16, 17]. Furthermore, heterogeneity is observed in direct (D1) and indirect pathways (D2) in the TS, which may further facilitate avoidance behaviour in threat-reward conflicts [18]. On the other hand, Greenstreet et al. [26] posit that TS-projecting neurons encode action prediction errors (APEs), which have been linked to “value-free” habits [33] and are difficult to integrate naturally into a standard value-based TDRL framework, as they are often modelled as a separate ad-hoc component [33, 34, 35]. The action prediction error hypothesis is further backed by experimental evidence showing neurons in the lateral substantia nigra pars compacta (SNc) and SNl that project to the TS respond to movement [22, 23, 24, 25], and Greenstreet et al. [26] raise the possibility that movement-related dopamine activity throughout the striatum may reflect action prediction errors. Reconciling these conflicting views is a central problem. Recent unifying accounts, for instance, highlight the TS’s role in stimulus-associated salience predictions but tend to disregard APEs [36]. Thus, a parsimonious account that integrates the normative roles of both TPEs and APEs into the TDRL framework remains elusive.

To address this, we adopt a normative framework from control engineering [37, 38, 39, 40], also known as linear RL [41] or entropy-regularised RL [42]. We propose that the dopamine system’s objective is not merely to optimise cumulative reward, but to optimise returns augmented by a penalty for deviating from a default policy. This single modification is useful for the present problem for two reasons. First, it provides a principled solution to the problem of optimally composing multiple value functions, for example corresponding to different reward or punishment types [43, 44, 45]. Second, it naturally introduces a default policy, which can be interpreted as a soft-habit policy [41]. In the TS model developed below, this default policy is updated in a value-free manner using action prediction errors [26, 33], but contributes to the valuation of future outcomes in a stable, off-policy manner [41, 45].

In this paper, we first briefly present the aspects of the entropy-regularised RL framework needed to model TS dopamine. Building on this, we present an instantiation of this framework—a computational model of TS-projecting dopamine signals—which suggests why aversive/threat prediction errors and action prediction errors may need to coexist. We then show that the model qualitatively reproduces key response patterns and simulations from previous studies [16, 17, 26]. Beyond this descriptive reconciliation, our model simulations highlight a normative role for TS-related computations in cautious behaviour under potential threat and stable learning under outcome uncertainty.

## 2 Results

### 2.1 Overview of the entropy regularised RL framework

This section provides an overview of the entropy regularised RL framework and presents it in comparison with the standard TD-RL model of phasic dopamine [1].

At Marr’s computational level [46], the standard TD-RL model optimises returns, i.e., cumulative discounted future rewards (equation 1.3). In contrast, the entropy-regularised RL framework formalises dopamine’s objective as maximising future returns augmented by a penalty for deviating from a default policy. The agent thus optimises a relative-entropy-regularised return (equation 1.11), where *KL*_*t*_ is the KL divergence between the current behavioural policy *π* and a default policy *π*^*d*^ at timestep *t* (equation 1.7). If *π*^*d*^ is uniform, this encourages random exploration (maximum-entropy RL); if *π*^*d*^ slowly tracks the learned policy *π*, it promotes choice perseveration or soft-habit formation by penalising deviations from recently taken actions [33, 41, 47, 48]. Lastly, we note that at the computational level, entropy-regularised RL is mathematically equivalent to planning/control as probabilistic inference [49].

At the algorithmic level, the standard model uses a temporal difference learning rule [2], with the temporal derivative of the striatal values computed and conveyed to the VTA to compute the RPE [1] (equation 1.1 and 1.2). The same algorithm can then be extended to state-action values (i.e., the SARSA algorithm) to drive action selection. The standard model has also been recently extended to multiple reward types [29], where a reward function is linearly decomposed into multiple rewarding attributes (*r*^*i*^), and individual values (*V* ^*i*^) optimised for each rewarding attribute are then linearly composed to result in scalar values used for action selection.

In the entropy regularised RL framework, we use an eligibility traces extension of soft Q-learning [50, 51, 42]. This is a temporal difference (TD (*λ*)) method, adapted here for the general entropy-regularised objective (equation 1.11). Soft Bellman state-value functions 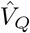 are computed from state-action values via a softmax weighted by the default policy (equation 1.4), making it off-policy, unlike the standard TDRL model or SARSA-based methods [1, 29]. Although the weighted softmaximum in equation 1.4 appears complicated, it is more biologically plausible than the hard maximum used in popular off-policy methods such as Q-learning, where computing max_*a*_ *Q*(*s, a*) is biologically intractable.

Because we use TD(*λ*) version of soft Q-learning (equation 43 in [50]) instead of soft Q-learning [42], the temporal difference error involves a biologically plausible term computed using the temporal derivative of the soft Bellman value function (equation 1.5, similar to equation 1.1) along with an additional KL divergence term (equation 1.6, 1.7). Alternatively, the same KL term also arises in the state value updates in the entropy regularised RL [50], but we utilise with state-action value updates here as they can directly drive behaviour in stochastic environments. In later sections, we demonstrate that this KL divergence term in the TD error presents clear similarities to the action prediction error (APE).

Importantly, similar to Millidge et al. [29], we can repeat the same learning process for multiple value functions 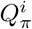 optimising the *i*-th rewarding attribute *r*^*i*^. These value functions can be composed using a weighted softmax (equation 1.8) to drive behaviour, using the same weights *w*^*i*^ applied to decompose the reward function (equation 1.9). The reward weights reflect current priorities, such as homeostatic needs [53, 29], inferred beliefs [27, 28], or other survival imperatives. Actions are then chosen using these composite Q-values via a Boltzmann policy (equation 1.10).

Mahajan and Seymour [45] demonstrate the following specific benefits of multi-attribute soft Q-learning over alternative approaches when applied to multi-attribute RL:

- Popular off-policy methods such as Q-learning, when extended to multi-attribute RL as in Dulberg et al. [54], can result in unreliable, sub-optimal composition; for example, the composite policy may treat two actions as equally preferable even when one action clearly optimises the composite reward function (see also [55]). In contrast, multi-attribute soft Q-learning guarantees reliable and optimal composition.
- Multi-attribute extensions of the standard TDRL model, such as Millidge et al. [29], can lead to unreliable learning under dynamically changing priority weights *w*^*i*^ due to their on-policy nature. Conversely, multi-attribute soft Q-learning guarantees reliable learning under changing priorities (reward weights).
- Multi-attribute soft Q-learning outperforms the successor representation [56], multi-attribute TD learning [29], and multi-attribute Q-learning [54, 55] in terms of fast adaptation to changing priorities.
- Multi-attribute soft Q-learning learns the same values as a compressed default representation (DR) [41] tuned to only relevant, predefined reward dimensions, recently also termed a terminal representation [57]. This offers two key benefits: it scales linearly, whereas the DR scales quadratically with the number of states, and it utilises more biologically plausible TD learning compared to the DR, which requires biologically intractable matrix inversions.

At Marr’s implementation level, the framework can account for heterogeneity in dopaminergic responses both between and within distinct neural targets. Separate value channels for distinct reward attributes (e.g., appetitive or aversive) support efficient multi-attribute learning while preventing positive outcomes from overriding threat or punishment values [58, 59, 60]. However, within a single dopaminergic target like the tail of the striatum (TS), we propose that heterogeneity can also arise from multiple value functions sharing a common outcome signal but differing in their initialisations, potentially reflecting priors for different contexts associated with different priorities. Such value initialisations can drive behaviour and explain dopaminergic responses even in the absence of explicit outcomes [32, 61, 17].

### 2.2 A computational model of TS dopamine

This section presents a particular instantiation of the aforementioned entropy-regularised RL framework: a computational model of the dopaminergic signals in the tail of the striatum (TS).

Several experimental findings inform the following key assumptions of our model:

- TS dopaminergic signals respond to aversive reinforcement (e.g., airpuffs) and not to appetitive reinforcement [15]. Therefore, the TS-relevant value functions utilise only the punishment signal (negative reward *ρ*, as per equation 2.5). In the absence of punishment, *ρ* = 0; otherwise, it is a negative signal.
- We propose that the TS-encoded Q-value function 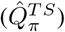 is composed of two Q-value functions corresponding to two different contexts: ‘Not-threatened’ 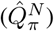 and ‘Threatened’ 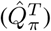, with different initialisations. 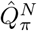 has a neutral (zero) value initialisation, whereas 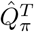 has a negative initialisation. Both value functions are updated separately using the shared *ρ* feedback as per equations 2.6 and 2.7. We later discuss how these two value functions with different initialisations may map onto direct and indirect pathways in the TS based on recent experimental findings by Tsutsui-Kimura et al. [18].
- The Q-value updates are scaled by both the learning rate *α* and eligibility traces *e*_*t*_(*s, a*) (equations 2.10 and 2.11). Equation 2.12 provides a simplified update rule for the eligibility traces applicable when using the soft-Bellman optimal policy (equation 2.19). Please refer to Mahajan and Seymour [51] for a minor variation involving importance sampling for action selection under any policy.
- The default policy is encoded by the TS alongside the two value functions and is updated using action prediction errors (APEs) as defined by Greenstreet et al. [26] and Miller et al. [33], which are equivalent to the updates used in Piray and Daw [41] (equation 2.9). The KL divergence between the behavioural policy and the default policy (equation 2.8) forms an additional term in the temporal difference errors used for updating the value functions (equations 2.6 and 2.7). We later demonstrate how this KL term also presents APE-like characteristics (equation 2.20), while naturally emerging in value updates (unlike the APE used in value-free updates of the default policy). The co-existence of both values and the default policy in the TS, alongside aversive and action prediction errors in TS-projecting SNl neurons, may be further supported by evidence that threat- and movement-related activity are encoded by separate dopaminergic subpopulations. Recent reports suggest threat- and acceleration-related dopamine responses are encoded in separate genetic subpopulations that both project to the TS, expressing Slc17a6 (also known as Vglut2) and Anxa1, respectively [62, 63, 64, 65, 26].
- The cortex keeps track of the probabilistic ‘threat’ belief state (*b*) alongside the agent’s location-based state (*s*). This is a belief over how likely the agent is to be in the ‘Threatened’ context as opposed to the ‘Not-threatened’ context. This belief state directly weights the composition of 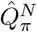 and 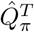 when computing 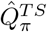 (equation 2.14). 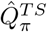 may be further composed with other reward-related striatal values using their corresponding reward weights (*w*^*R*^, equation 2.18) before driving behaviour (equation 2.19).
- Akiti et al. [17] developed the concept of Threat Prediction (TP, equivalent to the magnitude of an aversive value) as a value initialisation in a complete serial compound (CSC) model to explain experimentally observed TS dopamine responses. Similarly, Greenstreet et al. [26] utilise Action Prediction Errors (APEs) to explain TS dopamine responses in their data. We map the quantities in our model directly to the constructs used by Akiti et al. [17] and Greenstreet et al. [26]. We note that the mapping to threat prediction (TP) and the aversive prediction error with an APE-like KL term (equation 2.20) involves a negation, as values are learnt with negative feedback *ρ*, whereas TS-related dopamine signals encode the absolute magnitude of the aversive predictions and related errors. Therefore, our model (derived from the framework in Fig. 1) is a computational and algorithmic one rather than an implementation-level model. We acknowledge that excitation and inhibitory connections in *δ*^*N*^ and *δ*^*T*^ may potentially be reversed at the implementation level.

**Figure 1:**
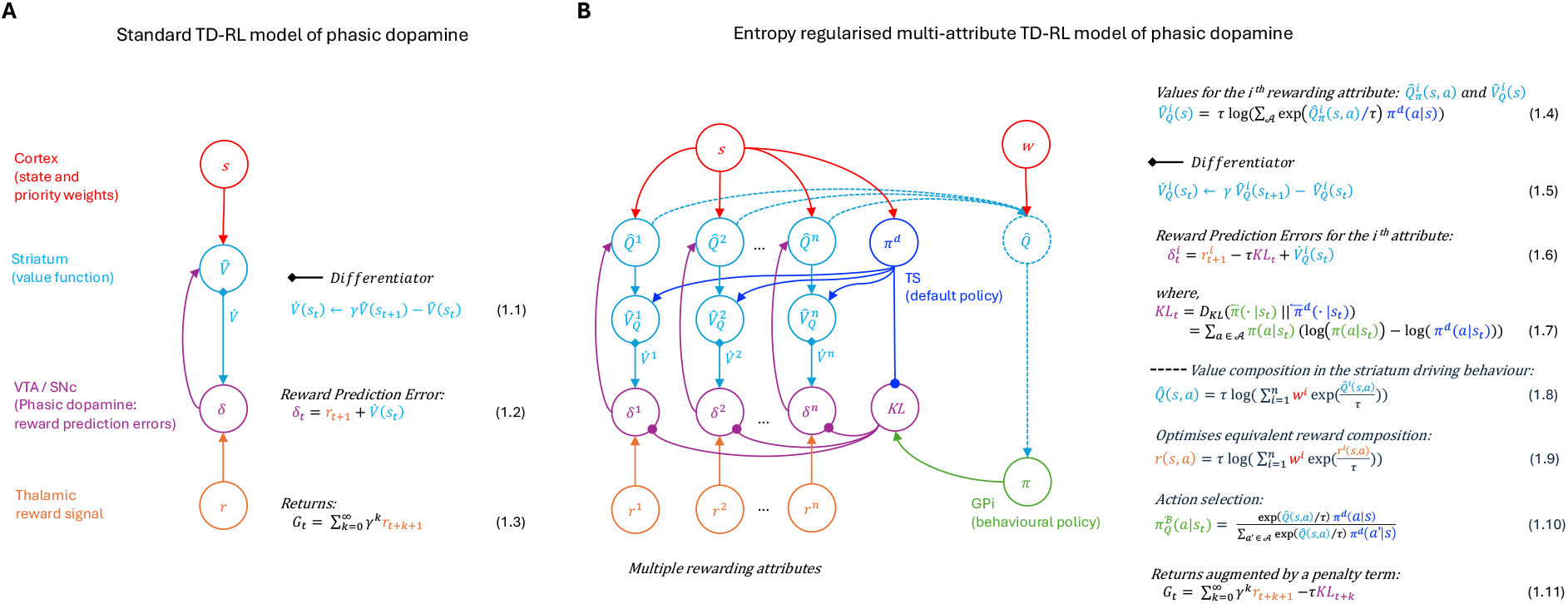
Overview of the standard TD-RL and proposed entropy-regularised RL framework of phasic dopamine. (A) The standard TD-RL model by Schultz et al. [1] involves computing the temporal difference error *δ*_*t*_ at time step *t* using the temporal derivative of the value function with discount factor *γ* (equation 1.1) and the reward feedback signal *r*_*t*+1_. The resultant algorithm optimises returns *G*_*t*_, defined as the sum of discounted future rewards (equation 1.3), following all conventions from Sutton et al. [52]. (B) Our proposed entropy-regularised framework provides a multi-attribute TD-RL model of phasic dopamine. It can be extended to optimise multiple rewarding attributes in parallel (*r*^*i*^), with *δ*^*i*^ explaining the DA heterogeneity pertaining to different reward types. All of the parallelly computed Q-values 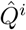 are combined using optimal weighted softmax-based composition (equation 1.8), leveraging the theory on the compositionality of optimal control laws [43]. The soft Bellman value function is computed based on the default policy *π*^*d*^ and not the behavioural policy, making it off-policy, unlike the standard TD-RL model [1] which is on-policy. Here, we utilise an eligibility traces extension to soft Q-learning [50, 51] which employs a temporal derivative in the soft Bellman value function (equation 1.5), similar to the standard model in panel A. The temporal difference error includes a KL term dependent on the behavioural policy and the default policy, in addition to the reward prediction error terms. At the computational level, the model optimises the returns augmented by a KL-penalty term (equation 1.11).

### 2.3 Simulations on threat prediction signals in TS dopamine during retreat from a novel object

Studies in mice reveal that interactions with a novel object involve approach-retreat bouts (Fig. 3A), which are importantly accompanied by TS dopamine activity during retreat but not during approach [16, 17] (Figs. 3E,F). Akiti et al. [17] model TS dopamine using ‘Threat Prediction’, i.e., the magnitude of the aversive value function. They model this phenomenon using a complete serial compound (CSC) model that incorporates value initialisation, equivalent to a potential-based reward shaping bonus [66]. The CSC model, albeit tremendously helpful as a descriptive qualitative model of the TS responses, has the following shortcomings:

- Since the CSC model assigns a separate state for each timestep rather than the agent’s location, the model can produce TS activity during retreat only if the aversive value initialisation is applied to states after the predefined retreat timestep. It is unclear how this kind of value initialisation can be extended to 2D state-action spaces.
- The CSC model cannot produce approach-retreat bout behaviours, as it is not designed for 2D state-action spaces (e.g., [67]).
- The CSC model relies on arbitrary thresholds for engage-avoid decisions.
- A shaping bonus is a non-distorting bonus [67], which would mandate that the net effect of any cycle of states (such as an approach-retreat bout) on the sum of prediction errors is zero. However, this condition cannot be sufficiently tested in a CSC model, which is unidirectional in time with no cycles.

To simulate the TS dopamine activity observed during retreat from the novel object, we designed a gridworld environment similar to the mice environment in Akiti et al. [17] (Fig. 3B). The grey grids in the vicinity of the novel object (purple grid) present a high probability (95%) of generating a threatening observation *o*_*t*_ = 1, whereas white grids in the rest of the gridworld present a high probability (95%) of generating a non-threatening observation *o*_*t*_ = 0. The time step when the agent is closest to the novel object and inside the grey grids is recorded as the start of the retreat. Our proposed model, as per Fig. 2, uses a belief state (*b*) over whether the underlying context is ‘Threatened’ or ‘Not-threatened’ to titrate the influence of the two value functions 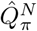 and 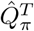. This belief state filter is implemented as equations 3.1–3.3 (Fig. 3C). 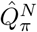 is initialised with zero values and 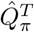 is initialised with a negative initialisation centred at the novel object, plotted in Fig. 3D. In addition to our proposed model, we also compare with a baseline model using only 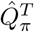 (i.e., *b* = 1), which is the same as a vanilla value initialisation in space, equivalent to potential-based reward shaping in the gridworld [67, 66]. For model simplicity, we do not include other reward-based value functions, but these can be included in the value composition before action selection.

**Figure 2:**
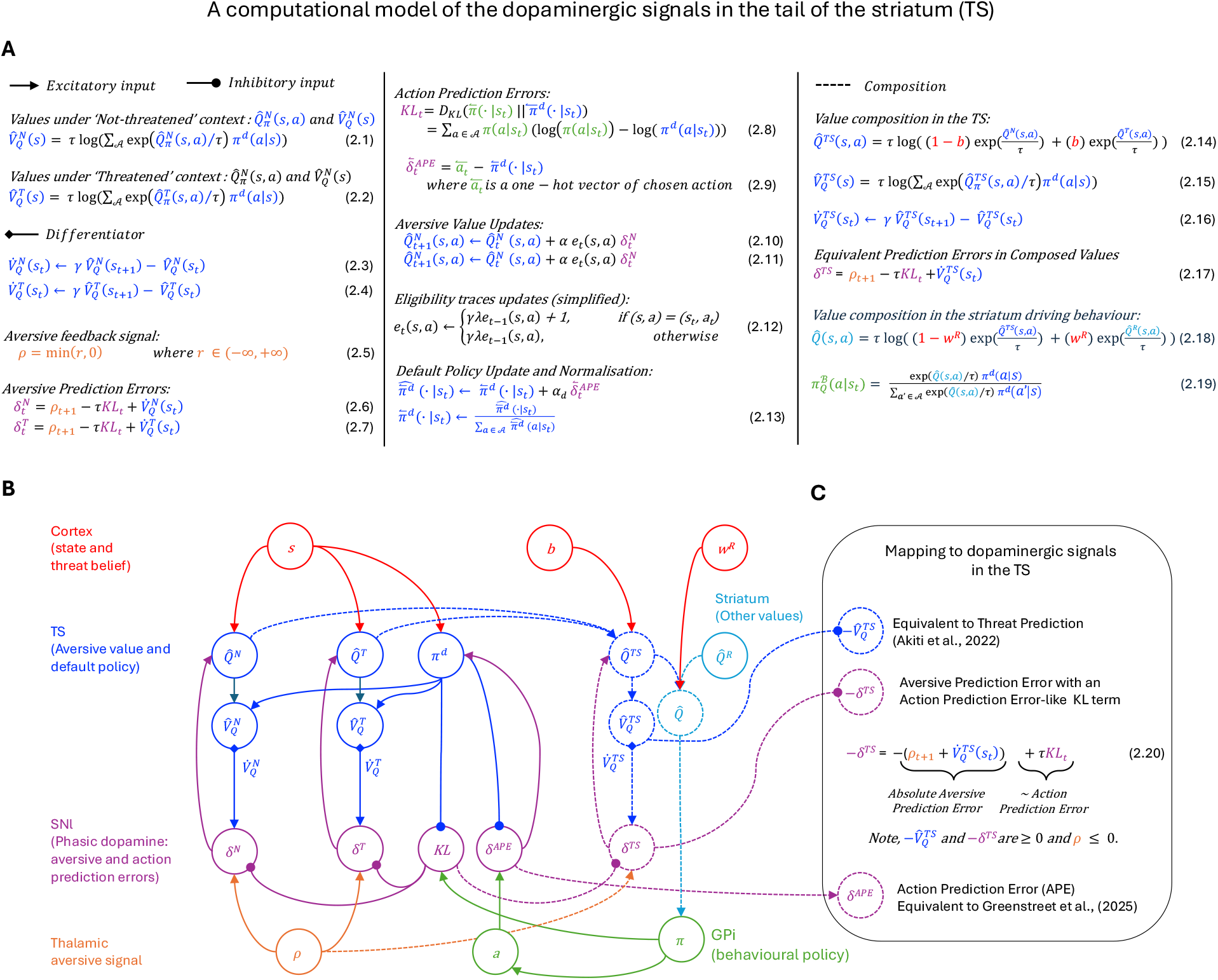
A computational model of the dopaminergic signals in the tail of the striatum. (A) Equations governing the proposed model. 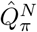 and 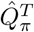 are Q-value functions for the ‘Not-threatened’ and ‘Threatened’ contexts, respectively, while 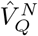 and 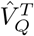 are the corresponding soft Bellman value functions. *ρ* is the aversive feedback signal (equation 2.5), shared between the value functions in both contexts, contributing to the aversive TD errors 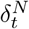 and 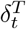. The default policy is updated based on the chosen action using action prediction errors, scaled by the learning rate *α*^*d*^, and then normalised (equations 2.9, 2.13). The aversive values are updated by scaling the aversive TD errors by the learning rate *α* and eligibility traces *e*_*t*_(*s, a*), which are updated as per equation 2.12. The TS Q-values are composed using weights derived from a belief state *b*, and can be further integrated with other reward-based value functions according to their respective weights *w*^*R*^. (B) A colour-coded wiring diagram of the proposed TS model. (C) The mapping of model signals to the dopaminergic signals observed in experimental evidence, and to previous models that attempt to explain them [17, 26]. Equation 2.20 further suggests why an aversive prediction error and an action prediction error-like KL term may co-exist in the TD errors, alongside previously proposed threat prediction (TP) values and APEs.

**Figure 3:**
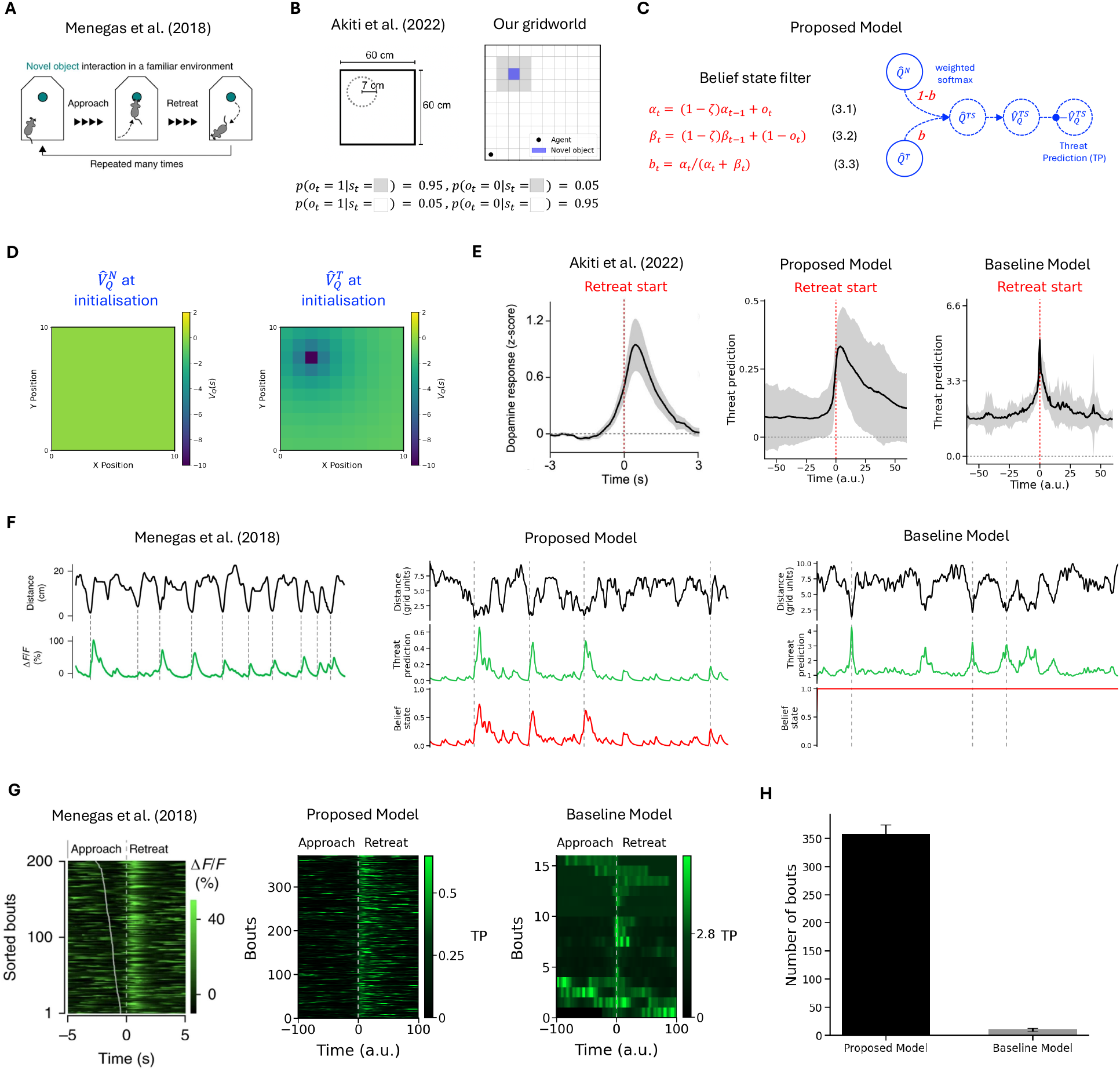
Simulation results on threat prediction (TP) signals in TS dopamine during retreat from a novel object. (A) Mice interacting with a novel object in a familiar environment demonstrated approach-retreat bouts [16]. (B) Our gridworld mimics the environment used by Akiti et al. [17]. The purple grid represents the novel object, and the threatening observations *o*_*t*_ are sampled from a Bernoulli distribution using the provided likelihood equations (for grey and white grids). (C) The proposed model involves inferring a belief state over whether the context is ‘Threatened’ or ‘Not-threatened’, and the posterior distribution is modelled using a Beta distribution with parameters *α*_*t*_ and *β*_*t*_. These parameters are updated iteratively based on the observation *o*_*t*_ with a decay term *ζ* = 0.1 (equations 3.1–3.3). (D) The proposed model utilises two different value initialisations: neutral for 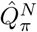 and aversive for 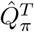. (E) Threat Prediction in the proposed model qualitatively reproduces the TS activity observed after the start of a retreat but not during approach [17], while the baseline model using only 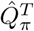 fails to do so. (F) The same result is plotted along a continuous timeline before averaging, alongside the data from Menegas et al. [16]. Δ*F/F* represents the TS activity. The proposed model demonstrates higher TP signals at the start of a retreat (denoted by a dotted line), unlike the baseline model. (G) The same result is plotted by stacking all bouts along the y-axis as a heatmap. Menegas et al. [16] sort the bouts by the start of the approach, while we do not. Our model shows higher values after retreat start (after the dotted line at time = 0), whereas the baseline model demonstrates high values before and after the retreat start. (H) The proposed model produces an order of magnitude more approach-retreat bouts than the baseline model. All figures are retrieved and minimally adapted from Menegas et al. [16] with permission from Springer Nature and from Akiti et al. [17] under CC-BY-NC-ND 4.0 license with permission from Elsevier.

We find that our proposed model qualitatively produces the key TS activity observed after the start of a retreat but not during approach [17] (Fig. 3E). Meanwhile, the baseline model fails to produce these temporally asymmetric signals, instead creating roughly equal activity before and after the retreat start (Fig. 3E). We demonstrate the same finding in Figs. 3F & G before averaging over the approach-retreat bouts, when compared to continuous data from Menegas et al. [16]. Additionally, since our model is operationalised over a 2D state-action space, we find it also qualitatively produces the observed approach-retreat bouts seen in experimental data (in distance to the novel object, Fig. 3F), and we find that the proposed model produced an order of magnitude more bouts than the baseline model, which cannot dynamically titrate the influence of 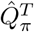 (Fig. 3H).

We do not include explicit curiosity-based exploration rewards for simplicity of modelling TS responses. However, their inclusion in the behavioural model could further increase agent tendencies to approach the novel object and make the distance to the novel object time-series less noisy, as currently the agent approaches the novel object through a series of random action selections rather than an intrinsic motivation to seek it. We observe in Fig. 3F that a majority of the Threat Prediction signals are governed by the belief state dynamics; however, we found in our additional experiments that an alternative non-Bayesian, switch-like dynamics in weights modulating the two values (Supplementary Fig. S1) also qualitatively produces similar TS responses during retreat, showing that the threat prediction response need not necessarily mimic the belief state response over time. Lastly, the magnitude of the TP responses in the proposed model are smaller than in the baseline model, as during weighted softmax composition of negative values, the exponent of negative values tends to be small. The same result can be qualitatively reproduced using a linear combination of the weights; however, we stick with weighted softmax due to the general benefits afforded by the compositionality of optimal control laws in our framework [43, 45].

At a theoretical level, the dynamic composition of multiple value initialisations as in the proposed model provides one way of unifying traditionally distorting novelty bonuses and non-distorting shaping bonuses [32], i.e., under fixed weights, it acts like a shaping bonus, and under varying weights, it produces novelty-like bonuses, all under a non-distorting scheme. In terms of testable predictions, we discuss in the Discussion section how the belief-gated, dual-value architecture offers a potential neural implementation for TS function, where 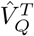 and 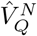 could potentially map onto direct (D1) and indirect (D2) TS pathways, respectively. Beyond that, our simulations suggest a hypothesis that gating an aversive value based on a threat belief state, akin to a context-dependent switch, may be a normative mechanism for adaptive safe exploration.

### 2.4 Simulations on action prediction errors in TS dopamine and their role in stable learning

While multiple value initialisations can account for threat prediction in the tail of the striatum (TS), recent findings also implicate TS dopamine in encoding action prediction errors (APEs) [26] that support perseverative behaviours or soft habits [33, 34, 47, 41]. This raises the question of how these seemingly distinct TPE and APE signals can be reconciled within a unified normative framework, and what computational purpose such APE-like signals serve.

The entropy-regularised RL framework offers a parsimonious explanation. The default policy *π*^*d*^, so far assumed uniform, can be slowly updated using action prediction errors as defined by Greenstreet et al. [26] and Miller et al. [33] to track the agent’s learned policy *π*, thereby encouraging perseveration [41]. The task performed by the mice in Greenstreet et al. [26] was a simple two-choice task based on a learned ability to distinguish between two contexts (high- and low-frequency tones). That study found that TS activity was higher in earlier trials and reduced in late trials as soft habits developed (Fig. 4A). To make the task slightly more challenging for simulation purposes, and to exploit the benefits of an eligibility-traces-based solution, we construct an analogous multi-step, two-choice task with different reward contingencies based on a fully observable context. The agent starts in state *S*_0_ and can choose either left or right actions to reach a goal state five steps away on either side. The agent has a higher probability of reward at the left goal in the high-frequency context, and vice versa (Fig. 4B).

**Figure 4:**
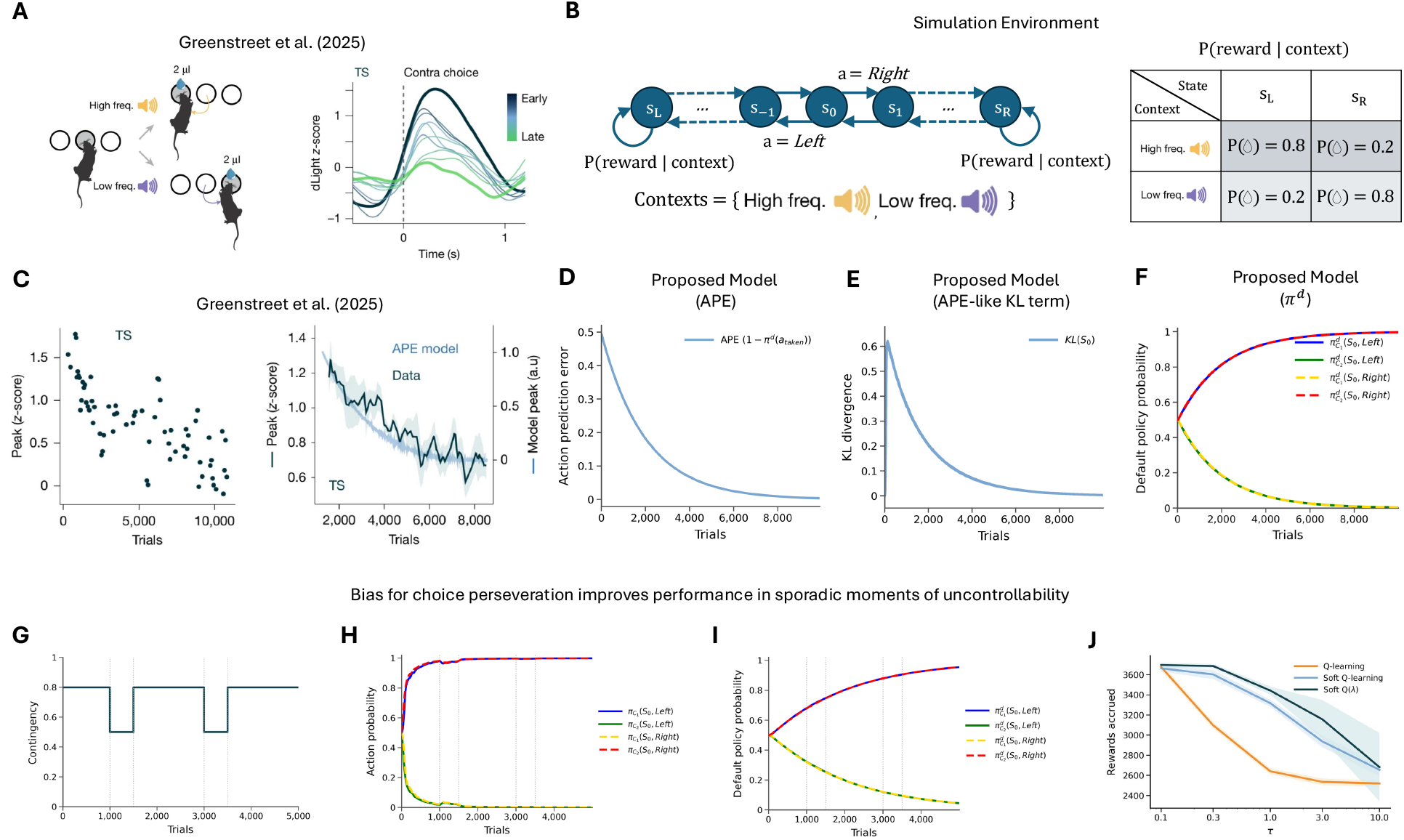
Simulation results on action prediction errors in TS dopamine and their role in stable learning. (A) The two-choice task paradigm from Greenstreet et al. [26], where subjects have to distinguish high- and low-frequency tones and make the correct choice to receive a reward. A left action in the high-frequency context results in a reward, and vice versa. The main experimental results show a decrease in TS activity as soft habits are acquired across the task. (B) Analogous multi-step probabilistic two-choice task with two different, fully observable contexts (high and low frequency). The agent starts at *S*_0_, and the goal states *S*_*L*_ and *S*_*R*_ are five steps away from *S*_0_. The reward contingency based on the context is provided in the table *P* (reward | context). (C) Greenstreet et al. [26] observe that TS activity decreases across trials as mice acquire soft habits, which they qualitatively model using action prediction errors (APEs) that drive “value-free” habits [33]. (D) The APEs used to update the default policy in our model (also in a value-free manner) reproduce the APE curves from Greenstreet et al. [26]. (E) The KL term in our proposed model initially shows an increase and then an APE-like decrease after a few early trials. (F) Default policy trajectories show the acquisition of soft habits across the task. (G) Task variant involving sporadic contingency degradation to test the role of the perseverative bias. (H) Evolution of the behavioural policy shows stickiness in actions. (I) Evolution of the default policy. Both panels H and I are plotted for the model with *τ* = 0.3. (J) Soft Q-learning and its TD(*λ*) extension outperform standard Q-learning across a range of *τ* values. All simulation plots are averaged over 10 runs. Figures adapted from Greenstreet et al. [26] under a CC-BY 4.0 license (figures were cropped and combined).

We find that the APEs used to update the default policy in our proposed model match the APE responses that Greenstreet et al. [26] used to qualitatively model their experimental data (Fig. 4C, D). Additionally, we observe that the KL term in the TD error starts at 0, as we initialise both *π* and *π*^*d*^ as uniform random policies; however, once it quickly rises in early trials, it demonstrates the same decrease observed in the APE signals (Fig. 4E). As the task involves no punishment, equation 2.20 results in only the *τ KL*_*t*_ term remaining, which we demonstrate shares clear APE-like patterns. This decrease is also reflected in the default policy *π*^*d*^ gradually aligning with the learned contextual policies *π*, effectively encoding soft habits (Fig. 4F). This result offers a unifying perspective: TS dopamine could reflect a composite signal where TPEs dominate in threat-relevant contexts, while an APE-like KL term dominates during the acquisition of soft habits.

Lastly, to understand the normative benefit of such APE-driven perseveration or stickiness in choices, we designed an experiment with the same simulation setup, except with sporadic moments of uncontrollability—specifically, for 500 trials following the 1000th and 3000th trials, where the reward contingency degraded from 0.8 to 0.5 (chance level; see Fig. 4G). We hypothesised that the bias towards *π*^*d*^ (soft habits) promotes learning stability during such contingency degradation. We found that during these uncontrollable periods, agents using soft Q-learning and its TD(*λ*) extension exhibited perseveration of prior choices and limited unlearning of the behavioural policy due to the influence of the slowly updated default policy (Fig. 4H, I). This resulted in better overall reward accrual (i.e., performance) compared to standard Q-learning (which lacks this perseverative bias) across a range of *τ* values (Fig. 4J). Therefore, we demonstrate that the KL penalty terms in the entropy-regularised return objective confer a value-based behavioural bias towards the default policy, which improves performance amidst sporadic moments of uncertainty or uncontrollability.

## 3 Discussion

There is considerable debate regarding whether dopamine in the tail of the striatum (TS) encodes action prediction errors (APEs) or aversive prediction errors. This debate has generated significant interest, with evidence supporting both sides [15, 16, 17, 36, 18, 26]. This paper presents a model that reconciles these conflicting views, explaining why both signals may need to coexist and qualitatively replicating the experimental data previously used to support each side. Beyond this descriptive reconciliation, our simulations highlight the normative role of the TS in cautious behaviours during potential threats, and in stable learning during outcome uncertainty.

The core contribution of this work is the proposal that midbrain dopamine, at a computational level, optimises cumulative rewards augmented by a penalty for deviating from a default behavioural policy. Applying this framework to the TS yields clear normative benefits: it dynamically titrates the influence of different value initialisations for safe exploration, and biases the behavioural policy to prevent deviations from the default policy—yielding performance improvements during sporadic contingency degradation. Furthermore, the flexible expression of innate values may reconcile associative and non-associative fear conditioning accounts [68], promoting more adaptable safe behaviour than previous models based solely on outcome uncertainty [59]. Lastly, our approach extends existing models [17] by demonstrating approach-retreat bouts in 2D state-action spaces and complement recent risk-sensitive model-based RL efforts in modelling cautious behaviours [69]. (Broader benefits of this framework for multi-attribute RL, off-policy learning, and reliable value composition are discussed at length in Mahajan and Seymour [45]).

Our framework synthesises several threads of the evolving dopamine story, which has progressively expanded from a simple scalar reward signal [1] to a multifaceted control signal. The model naturally accommodates aversive and threat prediction errors [14, 15, 16, 17], aligning with multi-threaded, outcome-specific prediction error views [13, 29]. This complements the feature-specific vector RPE model [30] while remaining compatible with it. Crucially, we propose an alternative account of within-target dopaminergic heterogeneity based on differentially initialised, composable values sharing a common outcome. We find that such value compositions unify novelty bonuses and (non-distorting) shaping bonuses in TDRL [32], which were previously used to explain early observations of novelty responses in phasic dopamine [70, 71].

Our proposal extends beyond the unifying hypothesis that the TS simply shifts attention to orient or avoid [36] by formally acknowledging action prediction errors and their normative implications for stable learning. Biologically, threat- and movement-related activity may be encoded by separate dopaminergic subpopulations. Recent reports support this, demonstrating that threat- and acceleration-related dopamine responses are encoded in distinct genetic subpopulations that both project to the TS [62, 63, 64, 65, 26].

Our work introduces the concept of an inferred threat belief state to titrate the balance between multiple values and guide flexible avoidance. This aligns with a performance effect, rather than a learning effect, in models of striatal direct (D1) and indirect (D2) pathway balance [31]. Mapping values 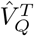 and 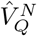 different initialisations to TS D1 and D2 pathways [18] respectively, generates several testable predictions:

1. **Opposing roles in conflict:** In a threat-reward conflict task, D1 pathways in the TS should promote avoidance from potential threats while D2 pathways in the TS should suppress it. This contrasts with the role of these pathways in rest of the basal ganglia, where D1 pathways usually promote approach to reward and D2 pathways suppress it [31].
2. **Ablation effects:** Ablating D1 TS neurons should reduce avoidance, whereas D2 TS ablation should increase it.
3. **Cue modulation:** If the inferred threat belief *b*_*t*_ covaries with observable threat cues (e.g., object size or movement), composite TS dopamine responses should scale accordingly.
4. **Distinct learning dynamics:** With repeated exposure to a novel object, the D1-associated 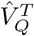 should diminish (reducing avoidance), while the D2-associated 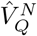 remains stable.
5. **Asymmetric TPEs:** In a one-step task, a punishing outcome with a magnitude between the initial 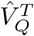 and 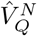 values should elicit negative TPEs in D1-projecting SNl neurons and positive TPEs in D2-projecting SNl neurons [16], potentially encoding outcome uncertainty following the computational proposal by Mikhael and Bogacz [72].

Recent work by Tsutsui-Kimura et al. [18] provides evidence supporting the first three predictions. Their findings on learning dynamics were mixed, and the prediction of asymmetric TPEs remains to be directly tested. Nevertheless, our model offers a robust explanation for within-target dopaminergic heterogeneity that is distinct from feature-specific accounts [30].

Finally, while the computation of APEs depends on the chosen action, the KL term in our model is action-independent and relies instead on the entire policy distribution in a state. We propose that this KL term provides a state-dependent analogue to APEs [26, 33]. Because it is action-independent, it easily scales to continuous actions—a challenge for traditional APE definitions. Furthermore, while APEs typically update the default policy in a value-free manner [33], the KL term actively updates the value function. This confers a value-based behavioural bias towards the default policy, providing stability during uncontrollable moments. As noted by Lindsey and Litwin-Kumar [35], Lindsey et al. [73], such mechanisms facilitate stable learning amidst motor noise and competition from distributed control systems. Lastly, our supplementary results present further connections to the model proposed by Bogacz [34] highlighting the role of dopamine in reward-based learning and action inference.

In conclusion, by proposing that the dopamine system optimises an entropy-regularised return, this paper provides a unified, normative account of TS dopamine. Our framework successfully reconciles threat and action prediction errors within a single computational model, demonstrating how the brain achieves safe, stable learning while generating clear predictions for future experimental inquiry.

## Author Contributions

PM: Conceptualization, Investigation, Formal Analysis, Software, Visualization, Writing – Original Draft Preparation, Writing – Review & Editing. BS: Funding Acquisition, Supervision, Writing – Review & Editing.

## Acknowledgments

An earlier extended abstract of this work was presented at RLDM 2025, and the authors thank the two anonymous reviewers at RLDM 2025, Thomas Akam and Chris Summerfield for their helpful suggestions on how to make this work more impactful. The authors thank Rafal Bogacz on very helpful suggestions that helped in rewriting and editing of the manuscript to making it more impactful, with a clear focus the debate on the tail of striatum, highlighting the canonical action prediction error in our model and on the presentation of the figures. Authors thank the funders: Wellcome Trust (214251/Z/18/Z, 203139/Z/16/Z and 203139/A/16/Z), IITP (MSIT 2019-0-01371) and JSPS (22H04998). This research was also partly supported by the NIHR Oxford Health Biomedical Research Centre (NIHR203316). The views expressed are those of the author(s) and not necessarily those of the NIHR or the Department of Health and Social Care. For the purpose of open access, the authors have applied a CC BY public copyright licence to any Author Accepted Manuscript version arising from this submission.

## Code and Data availability

Code for all simulations is available at https://github.com/PranavMahajan25/TSdopamine.git. No data was collected during this study.

## Supplementary Materials

### 3.1 Simulation parameters

**Table 1:** Summary of the hyperparameters used in simulation experiments.

| Hyperparameter | TP simulation | APE simulation | Choice Perseveration |
| --- | --- | --- | --- |
| State-space / layout | grid_size = 10 | steps_to_goal = 5 | steps_to_goal = 5 |
| Actions | up, down, left, right | Left, Right | Left, Right |
| Contexts | Threatened, Not Threatened | High Freq, Low Freq | High Freq, Low Freq |
| Reward / task setup | reward_positions at (7, 2) | fixed contingency task | contingency degradation |
| Contingency schedule | – | {0:0.8} | {0 : 0.8, 1000 : 0.5, 1500 : 0.8, 3000:0.5, 3500:0.8} |
| Discount factor $\gamma$ | 0.99 | 0.99 | 0.99 |
| Learning rate $\alpha$ | 0.1 (for both models) | 0.1 | 0.1 |
| Temperature $\tau$ | 0.5 | 0.1 | 0.3; sweep: 0.1, 0.3, 1.0, 3.0, 10.0 |
| Default-policy learning rate $\alpha^d$ | 0 | 0.001 | 0.001 (Soft Q and its TD($\lambda$) extension) |
| Trace parameter $\lambda$ | 0.5 | 0.5 | 0.5 |
| Initial Q-values | inverse-distance $\hat{Q}^T$ with max negative value = $-10$; $\hat{Q}^N = 0$ | all Q-values initialized to 0 | all Q-values initialized to 0 |
| Initial default policy | uniform: (0.25, 0.25, 0.25, 0.25) | uniform: (0.5, 0.5) per context | uniform: (0.5, 0.5) per context for Soft Q and its TD($\lambda$) extension |
| Belief-update parameters | decay $\zeta = 0.1$, threshold to determine end of retreat: $b < 0.1$ | – | – |
| Training horizon | max_steps = $10^5$ | num_episodes = 10000 | num_episodes = 5000 |

### 3.2 Relation to Bogacz [34]

In this section, we formally establish the connection between the Kullback-Leibler (KL) divergence penalty in our entropy regularised RL framework and the action prediction error (APE) derived in the Bayesian inference model of Bogacz [34].

We begin with the standard KL divergence between the current behavioural policy *π*(·|*s*_*t*_) and the default policy (or prior) *π*_*d*_(·|*s*_*t*_), which is defined as an expectation over the actions drawn from the behavioural policy:

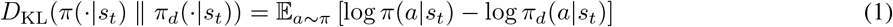

To relate this to Bogacz [34], we assume that both policies are parameterised as normal distributions with equal variance. Specifically, let the behavioural policy be *π*(*a* | *s*_*t*_) = *N* (*a*; *µ*^*π*^, *σ*^2^) and the default policy be *π*_*d*_(*a* | *s*_*t*_) = *N* (*a*; *µ*^*d*^, *σ*^2^), where *µ*^*d*^ is the mean of the default action prior in state *s*_*t*_.

Under these assumptions, we can decompose the KL divergence into two distinct components: a negative entropy term and an expected cross-entropy term:

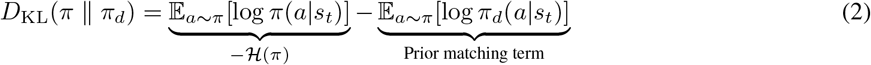

By substituting the Gaussian probability density function into the prior matching term, we obtain:

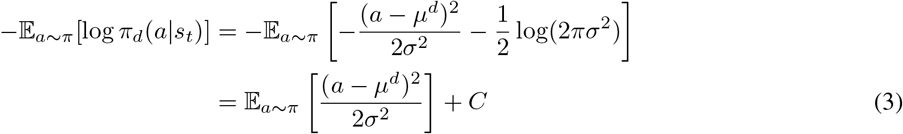

where *C* is a constant dependent on the variance. This formulation reveals that the KL divergence penalizes the expected squared deviation of the behavioural policy’s actions from the prior mean *µ*^*d*^, balanced against the entropy of the behavioural policy.

#### Sample-based Approximation

In practice, algorithms such as soft actor-critic (SAC) often rely on a single-sample Monte Carlo estimate of the KL penalty evaluated at the executed action *a*_*t*_ ∼ *π*(·|*s*_*t*_). Dropping the expectation yields the sampled penalty 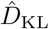:

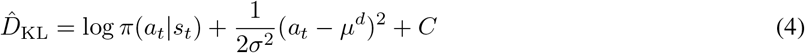

This single-sample approximation directly exposes the mathematical equivalence to the Bogacz [34] model. Bogacz [34] defines the habit prediction error (APE) simply as the deviation of the chosen action from the expected habitual action: *δ*_*h*_ = *a*_*t*_ − *µ*^*d*^.

Consequently, the sample-based KL penalty embedded within our regularized reward function incorporates the exact squared Action Prediction Error of the Bogacz model:

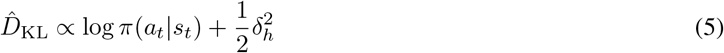

This demonstrates that penalizing the KL divergence from a passive prior in entropy regularised RL inherently applies a quadratic penalty based on the Bogacz [34] APE (*δ*_*h*_), alongside an entropy maximization bonus.

### 3.3 Additional results

**Figure S1:**
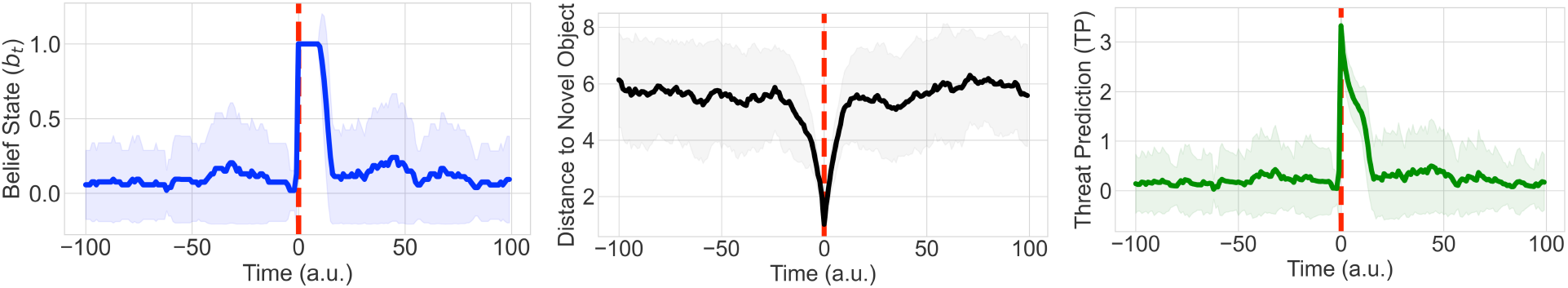
An additional experiment showing that gating the aversive initialisation with a belief state produces the desired temporal asymmetry in TP responses. (A) Belief state that acts like a switch turned on in the vicinity of the novel object and turned off randomly 10-20 steps after avoiding the object (B). Resultant distance to novel object showing the approach-retreat bout, and (C) The Threat-Prediction response on the retreat start, which decays along with the gradient of the value initialisation.

## Notes

### Competing Interest Statement

The authors have declared no competing interest.

